# A generic approach for studying the kinetics of liquid-liquid phase separation under near-native conditions

**DOI:** 10.1101/563700

**Authors:** Joris van Lindt, Anna Bratek-Skicki, Donya Pakravan, Ludo Van Den Bosch, Dominique Maes, Peter Tompa

**Affiliations:** VIB-VUB Center for Structural Biology, Vlaams Instituut voor Biotechnologie, Brussels, Belgium; Structural Biology Brussels, Vrije Universiteit Brussel, Brussels, Belgium; VIB, Center for Brain & Disease Research, Laboratory of Neurobiology, Leuven, Belgium; KU Leuven, Department of Neurosciences, Experimental Neurology, Leuven, Belgium; Institute of Enzymology, Research Centre for Natural Sciences of the Hungarian Academy of Sciences, Budapest, Hungary

**Author notes:** Joris van Lindt and Anna Bratek-Skicki contributed equally to the presented work.

## Abstract

Understanding the kinetics and underlying physicochemical forces of liquid-liquid phase separation (LLPS) is of paramount importance in cell biology, requiring reproducible methods for the analysis of often severely aggregation-prone proteins. Frequently applied approaches, such as dilution of the protein from an urea-containing solution or cleavage of its fused solubility tag, however, often lead to very different kinetic behaviors. Here we suggest that at extreme pH values even proteins such as the low-complexity domain (LCD) of hnRNPA2, TDP-43, and NUP-98 can be kept in solution, and then their LLPS can be induced by a jump to native pH, resulting in a system that can be easily controlled. This approach represents a generic method for studying LLPS under near native conditions, providing a platform for studying the phase-separation behavior of diverse proteins.

---

Compartmentalization is a basic device of eukaryotic cells for the regulation and spatiotemporal separation of their biochemical reactions. Many compartments termed organelles are surrounded by a membrane which physically separates them from the bulk cytoplasm. Cells, however, also contain many so-called “membraneless organelles” (MOs), which lack a physical barrier. MOs are involved in many cellular activities including metabolic processes and signaling pathways. Many studies suggest that liquid-liquid phase separation (LLPS) is responsible for the creation of these supramolecular assemblies.^1-3^ Recently, intense research has been focused on the influence of a prominent MO, stress granule, on cell survival and its link to neurogenerative diseases, such as amyotrophic lateral sclerosis (ALS).^4, 5^ Stress granule proteins form liquid droplets, then slowly undergo gelation and, in the end, are converted into aggregated fibrils. TDP 43, FUS and heterogeneous nuclear ribonucleoprotein A2/B1 (hnRNPA2/B1) are prime examples of phase-separating stress granule proteins. Similar transformation, from solution to characteristic “FG particles”, was also observed for the nuclear pore complex (NPC) protein NUP 98, which plays an important role in the bidirectional transport across the NPC.^6^ Due to its prevalence in a broad range of physiological and pathological processes of the cell, understanding the biophysical principles that govern LLPS is a central goal in current cell biology research.

Studies devoted to the phase separation phenomenon are dominated by the visualization of mature liquid droplets by a wide range of techniques, at a stage considered to correspond to equilibrium. Phase-separated droplets, however, are never in thermodynamic equilibrium but are in a transition towards a final state of two separate phases. Moreover, the proteins involved in these processes are aggregation prone and have a tendency to be sticky; thus, most often they are prepared either with a fused solubility tag (MBP, GFP, or GST),^7, 8^ or under denaturing (6-8 M urea)^9^ or otherwise non-physiological (very high salt, detergents)^10^ conditions, which may strongly interfere with experimental results. For example, transitions of the prion-like domain of Sup35 have been studied under two different conditions, starting from highly denaturing conditions (8M GuHCl) or from a stock of very high salt;^10^ from initial denaturing conditions, it aggregated into amyloid-like fibers,^11^ whereas when diluted from high salt, it phase separated into liquid droplets which later turned into gel condensates.^12^ It is, therefore, obvious that LLPS is very sensitive to experimental conditions which may lead to very different conclusions in terms of preferred trajectory along the transition path: solution → droplet → gel → amorphous or amyloid aggregate. In accord, the literature is full of qualitative statements about whether a protein phase separates under given conditions, without quantitative data on the underlying kinetics and parameters that significantly influence this phenomenon. Due to the general biological importance of the phenomenon of LLPS, the field requires clear experimental standards for developing physiologically relevant models based on reproducible results.^10, 13^

Toward this goal, we propose here a generic method in which phase separation is induced by a simple pH jump at near-native conditions. To demonstrate the benefits of our approach, we compared the kinetics of LLPS of the low-complexity domain (LCD) of hnRNPA2 initiated by a pH drop (from 11.0 to 7.5) with kinetics performed in the presence of salt, urea and by cleaving its MBP solubility tag as reported in literature^9^. We show that different conditions have a profound effect on kinetics of LLPS, which may lead to misinterpretation of obtained results. We also demonstrate that this principle provide a generic approach for studying the LLPS of practically any other aggregation-prone protein. By studying its pH-dependent net charge profile, an appropriate pH at which the protein stays in solution can be selected, and its LLPS studied, as demonstrated through the example of two other disordered proteins: the LCD of TDP 43 and a fragment of NUP98 (Fig1S, Supporting Information).

First, we registered absorbance changes at 600 nm (A600) of hnRNPA2 LCD solution after pH dropped from 11.0 (10 mM CAPS) to 7.5 (Fig. 1a, orange line). The turbidity of the solution increased rapidly reaching maximum value within minutes, which is attributed to the formation of droplets accompanying phase separation. To confirm the phase separation, we spun down solutions at pH 11.0 and 7.5 and using gel electrophoresis, we analyzed their supernatant and pellet. At pH 11.0 there was no pellet and the protein was present in the supernatant (Fig. 1a). However, at pH 7.5, the supernatant was almost protein-free while in the pellet most of the protein was detected, confirming that the protein was mostly present in the condensed form. An increase of ionic strength by 150 mM NaCl (Fig. 1a, blue line) slowed down the kinetics, indicating the importance of electrostatics in LLPS.

**Fig 1.**
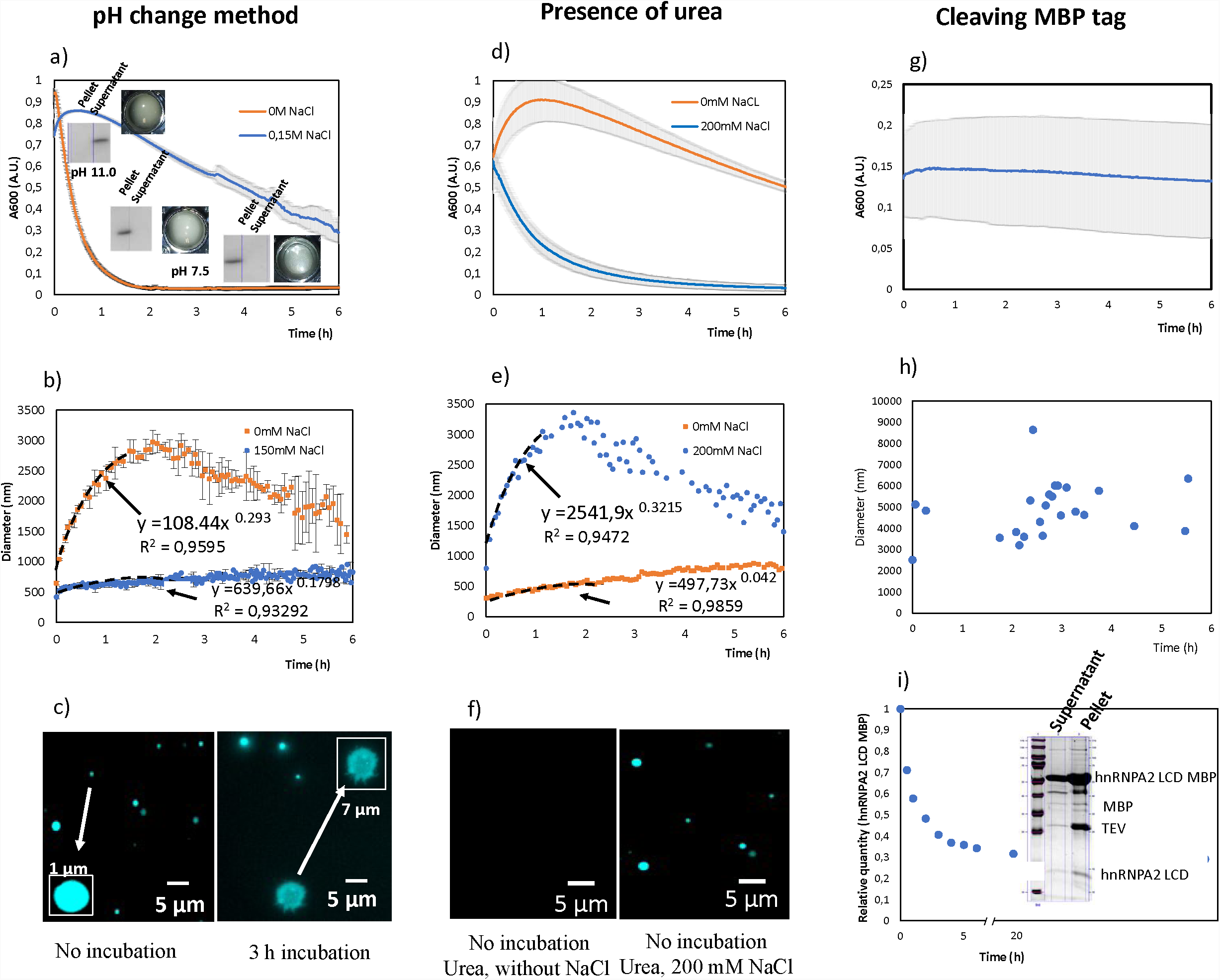
Phase separation (PS) of hnRNPA2 LCD initiated by three methods: i) by pH change (left side), ii) in the presence of urea (in the middle), iii) by cleaving the MBP tag (right side); a) PS of hnRNPA2 LCD monitored by turbidity without NaCl (orange line), with 150 mM NaCl (blue line) with inserted images of SDS PAGE gels representing the content of the supernatant and the pellet after incubation time of 1 min, 30 min, and 1h, b) size evolution of droplets monitored by DLS without NaCl (orange line), with 150 mM NaCl (blue line) c) fluorescence microscopy images of droplets without incubation and after 2 h incubation, d) PS of hnRNPA2 LCD in the presence of urea monitored by turbidity without NaCl (orange line), with 200 mM NaCl (blue line), e) size evolution of droplets monitored by DLS without NaCl (orange line), with 200 mM NaCl (blue line), f) fluorescence microscopy images of droplets formed in the presence of urea (left side) and with the presence of urea and 200 mM NaCl (right side),g) PS of hnRNPA2 LCD induced by cleaving the MBP tag monitored by turbidity, h) DLS measurements of hnRNPA2 LCD droplets formed after cleaving the MBP tag, i) cleavage rate of the hnRNPA2 LCD MBP protein with inserted image of SDS PAGE gel representing the content of the supernatant and the pellet after cleaving the MBP tag.

LLPS was also demonstrated by following the evolution of the droplets in time by dynamic light scattering (DLS) (Fig. 1b). Initially, small droplets of a diameter of about 1 μm were formed, growing slowly to a maximum of 3.0 μm in diameter after approximately 2 hours (Fig. 1, re line). During the increasing stage, the droplet radius increased in time with an exponential dependence of 1/3 (t~1/3), as expected for particle growth by Ostwald ripening.^14^ By about five hours after initiating phase separation, DLS measurements could no longer be interpreted, probably due to aggregation characterized by a high polydispersity. DLS also confirmed the importance of electrostatics, as in 150 mM NaCl it showed a significant decrease in droplet size, with a maximal value of 800 nm after 6 hours (Fig. 1., blue line). The evolution of droplets was also monitored by fluorescence microscopy (Fig. 1c). As DLS data suggests, small droplets were formed at the beginning of the experiment and grew over time. After 3 hours, approximately, large droplets were observed together with aggregates proving that the LLPS of hnRNPA2 LCD is not an equilibrium system.

The advantage of this approach for studying the kinetics of early stages of phase separation follows from that upon pH jump, there is no residual denaturant in the system (unlike upon dilution from 8M urea, for example), and it can be administered instantaneously by the addition of a small volume of concentrated buffer (unlike with the slow cleavage of the solubility tag). To demonstrate that these generally applied approaches can lead to potentially erroneous and artefactual results, we first initiated phase separation by diluting hnRNPA2 LCD stored in 8M urea into 20 mM HEPES, pH 7.4 buffer, resulting in a final urea concentration of 150 mM. The increase of turbidity is much slower, reaching maximum after 1.0 hour, then slowly decreasing over time (Fig. 1d, blue line). DLS data for the sample with urea confirmed the presence of particles (droplets) whose size at the beginning of the experiment was approximately 400 nm, then slowly increased reaching ca. 800 nm after 6 hours (Fig. 1e, orange line). Unlike with the pH-jump system, a fit of the curve at low salt gives an exponent of 0.1798, which is inconsistent with the process of Ostwald ripening. As an additional discrepancy, when we added 150 mM urea to the hnRNPA2 LCD sample kept at pH 11.0, and initiated phase separation by the pH jump to 7.5, we observed the same turbidity and DLS kinetics as without urea (Fig.1d, orange line and Fig.1e, blue line), suggesting that urea at a high concentration, strongly interacts with protein, influencing its path of LLPS.

When LLPS was induced by a pH jump in the presence of 150 mM urea and 200 mM NaCl, kinetics again followed a path observed by the pH change in the absence of salt (Fig. 1d, orange line vs Fig. 1a, orange line). DLS measurements also confirmed that the size evolution of the droplets was the same as observed by the pH jump method (Fig. 1e, blue line vs Fig. 1b, orange line). Previous research on protein-urea interactions have shown that urea is able to solvate protein backbone and sidechains, explaining urea denaturation,^15, 16^ which may be prevented by strong salt-urea interactions.^17-19^ The effect of residual urea in the presence of 200 mM NaCl was also demonstrated by fluorescence microscopy (Fig. 1f). We observed large droplets after diluting protein-urea sample in the presence of 200 mM NaCl while no droplets were observed without NaCl probably due to their small size as was confirmed by DLS.

We also demonstrated adverse effects on LLPS kinetics of hnRNPA2 LCD by cleaving the MBP solubility tag with TEV protease. A striking difference from the pH jump was observed, as at the given protease concentration, turbidity of the solution did not significantly change for a period of 6 hours, reaching an average value of 0.15 (Fig. 1g). The DLS data showed that the size of the droplets also remained constant having an average diameter of 5.0 μm without any indication of further transition(s) (Fig. 1h), i.e., the system apparently failed to demonstrate a behavior typical of LLPS. Therefore, we started a detailed analysis of the cleavage process. First, we ran aliquots of the phase separating solution taken at different times on SDS PAGE to follow cleavage of hnRNPA2 LCD-MBP. Interestingly, the cleavage was always incomplete, reaching plateau after a few hours (Fig. 1i). We hypothesized two explanations: i) either TEV got sequestered in the LLPS droplets, or ii) the hnRNPA2 LCD tail of hnRNPA2 LCD MBP was recruited in the LLPS droplet, protecting it from the cleavage. To prove our hypothesis, we spun down the droplets and analyzed the supernatant and the pellet via gel electrophoresis. We found, as expected, that most of hnRNPA2 LCD was present in the droplets (Fig. 1i). However, the most unexpected finding was the presence of TEV, the hnRNPA2 LCD MBP and the MBP tag in the droplets. Thus, in contrast to previous experiments, droplet content was very heterogenous. Therefore, experiments in which phase separation is induced by cleaving a soluble tag should be carefully analyzed.

These results have all shown that the pH jump system in the case of hnRNPA2 LCD enables to dissect its kinetics of LLPS under near-native conditions. Next, we sought to demonstrate that this approach can be applied to other proteins, by observing kinetics of LLPS of TDP-43 LCD and a fragment of NUP98 (Fig. 2). First, we have analyzed the net charge-pH curve of the two proteins, to select an appropriate pH where the proteins may not aggregate (Fig. 2a). We selected pH = 3.5 for both TDP-43 LCD and NUP-98 LCD. As appears, both proteins are in solution at the respective pH selected, and readily undergo LLPS when their pH changes to 7.5 and 10.0 for TDP 43 LCD and NUP 98 LCD, respectively. Phase separation was confirmed by turbidity assays (Fig. 2b, Fig. 2e), DLS measurements (Fig. 2c, Fig. 2f), which provided an exponent of 1/3 indicative of Ostwald ripening, and fluorescent microscopy (Fig. 2d), which showed the development of μm-sized droplets.

**Fig 2.**
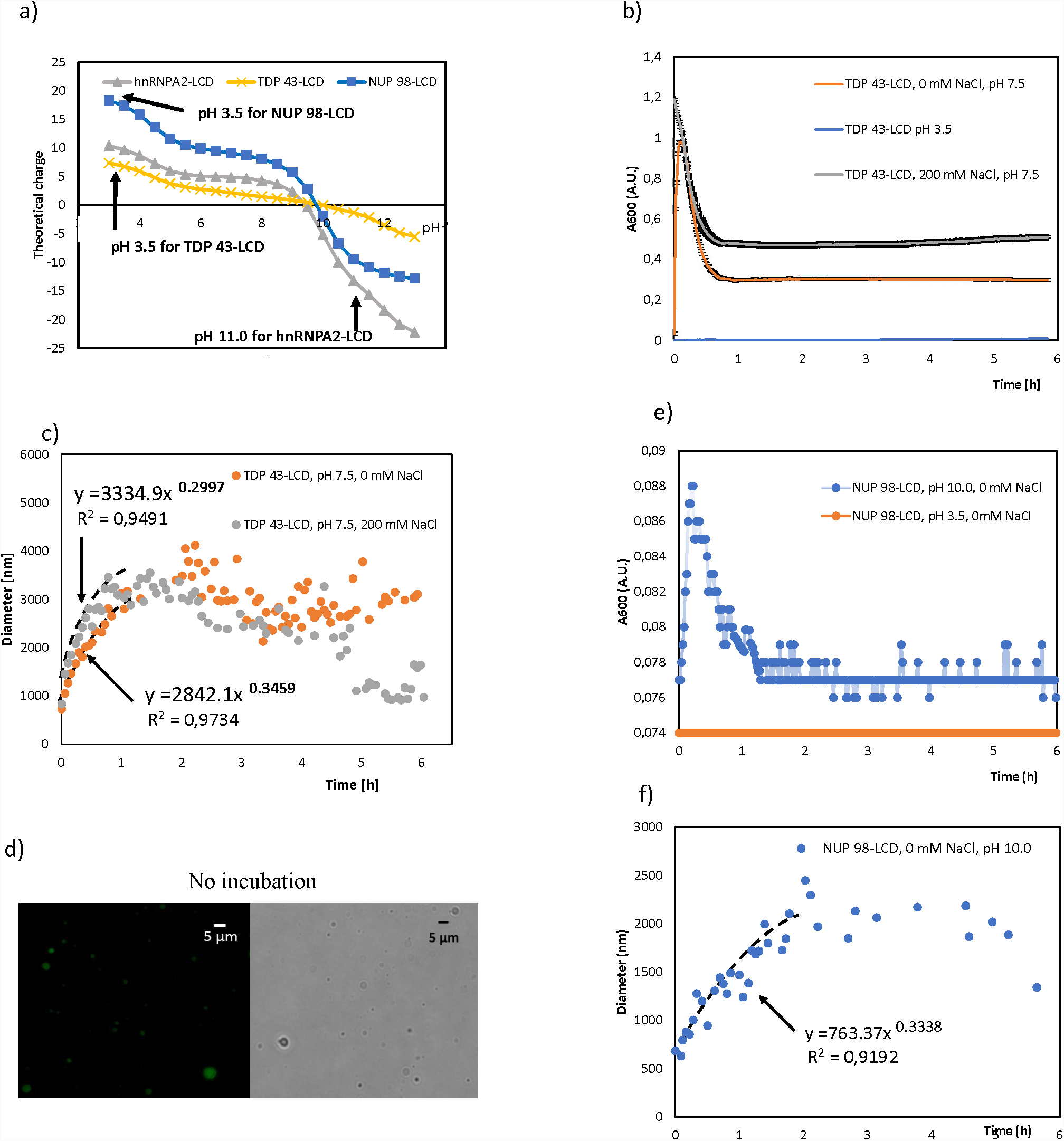
Phase separation of TDP 43-LCD and NUP 98-LCD: a) dependence of theoretical charge on pH of hnRNPA2-LCD (grey line), TDP 43-LCD (yellow line), NUP 98-LCD (blue line) with pH values at which proteins remain in solution, b) PS of TDP 43-LCD monitored by turbidity without NaCl (orange line), with 150 mM NaCl (gray line), c) size evolution of TDP 43-LCD droplets monitored by DLS without NaCl (orange dots), with 150 mM NaCl (gray dots), d) fluorescence and bright field microscopy images of droplets of TDP 43-LCD without incubation, e) PS of NUP 98-LCD monitored by turbidity without NaCl (blue dots), f) size evolution of NUP 98-LCD droplets monitored by DLS without NaCl (blue dots).

In conclusion, we present a method that can be used for studying the kinetics of early stages of LLPS of aggregation-prone proteins. By selecting an appropriate pH, the protein can be kept in solution and its phase separation can be initiated by a change of its pH. As the pH jump requires the addition of a small amount of concentrated buffer only, the composition of the system is free to vary and can be accurately controlled, thus providing a generic approach that is devoid of artefacts arising from the presence of residual denaturant and/or the slow and incomplete cleavage of a solubility tag.

## Acknowledgements

This work was supported by the Odysseus grant G.0029.12 from Research Foundation Flanders (FWO), a VUB Spearhead grant (SRP51, 2019-24) and grant K124670 from the Hungarian Scientific Research Fund (OTKA).

## Author Contributions

J.V.L, P.T., L.V.D.B, and A.B.S. conceived experiments and analyzed data. J.V.L, and A.B.S. carried out experiments, D.P. performed microscopy measurements. D.M. contributed to the initial design of the experiments. A.B.S. and P.T. wrote the manuscript and all authors were involved in revising it critically for important intellectual content.

## Materials and Methods

### hnRNPA2 LCD expression and purification

N-terminally polyhistidine-tagged hnRNPA2 LCD gene (R190 – Y341; UniProt P22626), located on a prokaryotic expression vector plasmid pJ411 with a kanamycin resistance gene as a selectable marker, with the lacI gene, and preceded by lac operon was a gift from Prof. Dr. N. Fawzy.

*E. coli* BL21 STAR (for protein expression), or NEB5α (for plasmid purification) were heat transformed in LB medium at 42°C. After 30 minutes at 37°C, kanamycin was added, and the solution was incubated overnight and shaken at 180 rpm. From this solution, a single colony of *E. coli* cells were obtained by spreading on agar plates with kanamycin, from which a glycerol stock was prepared and stored at −80°C.

Precultures were prepared by dipping a sterile toothpick in the glycerol stock and incubating it in 50 ml LB with kanamycin. After overnight incubation, the precultures were added to 1 L of terrific broth medium and incubated until they reached OD of 0.6 to 0.8 at 37°C and 170 rpm. 1mM Isopropyl β-D-1-thiogalactopyranoside was added to induce hnRNPA2 LCD expression, and the temperature was lowered to 26°C. After overnight incubation, cells were harvested by centrifugation at 4°C, 5000 rpm for 20 minutes. The cells were flash frozen and stored at −80°C.

The cells were thawed and resuspended in lysis buffer: 20 mM Tris-Cl, 500 mM NaCl, 10 mM imidazole, 1mM dithiothreitol supplemented with 0.1mM phenylmethylsulfonyl fluoride (PMSF), 0.5 mM benzamidine hydrochloride (BA), and 1 tablet Roche complete EDTA-free protease inhibitor per 50 ml. The cells were lysed by sonication (on a Sonics VCX-70 Vibra cell) for 15 minutes (5 seconds pulse on, 5 seconds pulse off, 70% amplification) on ice to avoid heating the sample. Inclusion bodies, containing hnRNPA2 LCD, were pelleted by centrifugation at 24 000 x g for 1h at 4°C. The pellet was resolubilized in a denaturing buffer: 20 mM Tris-Cl, 500 mM NaCl, 10 mM imidazole, 1 mM DTT, 3 M urea, pH 8.0 and centrifuged for 1 h at 24000 x g and 4°C to pellet bacterial debris. The supernatant was filtered through a filter of 0.45 μm and loaded onto a nickel-charged IMAC column (HistrapTM HP – GE Healthcare), to which the polyhistidine tag of hnRNPA2 LCD binds. Bacterial debris was washed away with a denaturing buffer, after which the bound hnRNPA2 LCD was eluted with a linear 0 mM to 250 mM imidazole gradient.

The eluted hnRNPA2 LCD with polyHis tag was cleaved by polyHis-tagged TEV protease in a buffer of 50 mM NaH_2_PO_3_, 20 mM NaCl, 3 M urea, pH 7.0 at room temperature. After overnight incubation, the cleaved tag and the protease were removed by running the solution over a nickel-charged IMAC column, collecting the flow-through containing the cleaved protein.

Purified hnRNPA2 LCD was extensively dialyzed: 2 x 4 h, once overnight in a 0.01 M CAPS buffer, pH 11.0 applied at a 1:100 concentration ratio. The protein was stored at a final concentration of 20 μM at – 80°C. Protein concentration was determined by densitometry of Coomassie-stained SDS-PAGE gels, and by QUBIT®.

### TDP 43 LCD expression and purification

60 μl of *E. coli* BL21 STAR cells were transfected with 1 ul of TDP43 LCD plasmid and incubated overnight (16 h) in 50 ml LB medium with 50 μl of LB medium with 50 ul of kanamycin (100 mg/ml) at 37°C. Next, 8 ml of the cells grown overnight were injected to 1 L of NZYM containing 1 ml of kanamycin and incubated at 37°C with continuous shaking at 170 rpm. After, approximately, 3 h when the OD was higher than 0.6, 500 ul of IPTG was added into the solution and the cells grew for the next 4 hours to be optimally expressed (OD ~ 2.0). The cells were harvested by centrifugation at 4°C, 5000 rpm for 15 minutes. A pellet from each 1 L solution was transferred into 1 falcon for maximal protein retention in the purification procedure. The cells were immediately frozen and stored at −80°C.

In the next step, the cells were thawed and resuspended in lysis buffer: 500 mM NaCl, 20 mM Tris HCl, 1 mM DTT, 1 tablet Roche complete EDTA-free protease inhibitor, 0.5 mM BA, 0.1 mM PMSF, pH 8.0 and sonicated in ice for 15 minutes at 70% amplitude applying 10 seconds on and 10 seconds off cycles.

Inclusion bodies were then pelleted by centrifugation at 5000 x g for 5 min at 4°C. The pellet was resolubilized in a denaturing buffer: 20 mM tris HCl, 8 M urea, 500 mM NaCl, 10 mM imidazole, 1mM DTT, pH 8.0 and sonicated in ice for 5 minutes or until the solution was clear as water (usually it was cycles of 5 seconds on and 5 seconds off at 70% amplitude). The solution was centrifuge at 20 000 x g for 45 min at 4°C. The supernatant was filtered through a filter of 0.45 μm and loaded onto a nickel-charged IMAC column (HistrapTM HP – GE Healthcare), to which the polyhistidine tag of TDP 43 LCD binds. Bacterial debris was washed away with a denaturing buffer after which the bound TDP 43 LCD was eluted with a linear 0 mM to 500 mM imidazole gradient. The protein fractions were combined, pH was adjusted to 7.0 and the following reagents were added: 1 tablet protease inhibitor, 0.1 mM PMSF, and 0.5 mM of BA.

Next, the solution was portioned in 1ml eppendorfs with 1/10 of TEV protease. All samples were incubated overnight at 34°C without shaking.

The protein samples were combined again, and the solution was loaded onto a HiPrep 26/10 desalting column using a 20 mM MES buffer, pH 5.5. The protein was filtered with 0.22 μm stored at a final concentration of 55 μM at – 80°C. The effectiveness of each purification step was determined by gel electrophoresis with Page Blue staining. Protein concentration was determined by a QUBIT® kit.

### NUP 98 expression and purification

The plasmid of Nup98 was provided by Addgene and expressed in E. coli BL21 cells. Protein in inclusion bodies was extracted and solubilized in a buffer of 8 M urea, 100 mM Na_2_HPO_4_, 10 mM DTT and 10 mM Tris-HCl, pH 8.5, purified through Ni-NTA affinity chromatography. The eluted protein was finally dialyzed into a PBS buffer pH 3.0 and kept frozen at –20°C in small aliquots.

### hnRNPA2 LCD MBP expression and purification

*E. coli* BL21 STAR were heat transformed in LB medium at 37°C. After 30 minutes at 37°C, kanamycin was added, and the solution was incubated overnight and shaken at 180 rpm. From this solution, a single colony of *E. coli* cells were obtained by spreading on agar plates with kanamycin, from which a glycerol stock was prepared and stored at −80°C.

Precultures were prepared by dipping a sterile toothpick in the glycerol stock and incubating it in 50 ml LB with kanamycin. After overnight incubation, the precultures were added to 1 L of terrific broth medium and incubated until they reached OD of 0.6 to 0.8 at 37°C and 170 rpm. 1mM Isopropyl β-D-1-thiogalactopyranoside was added to induce hnRNPA2 LCD MBP expression, and the temperature was lowered to 28°C. After incubating overnight, the cells were harvested by centrifugation at 4°C, 5000 rpm for 20 minutes. The cells were flash frozen and stored at −80°C.

The cells were thawed and resuspended in a lysis buffer: 100 mM KCl, 50 mM Tris-Cl, 1 M NaCl, 10 mM imidazole, 1mM dithiothreitol supplemented with 0.1mM PMSF, 0.5 mM BA, and 1 tablet Roche complete EDTA-free protease inhibitor per 50 ml. The cells were lysed by sonication (on a Sonics VCX-70 Vibra cell) for 10 minutes (5 seconds pulse on, 5 seconds pulse off; 70% amplification) on ice to avoid heating the sample. Then, the solution was incubated for 30 min at 37°C with 50 μg/ml DNase and 20 mM MgCl_2_.

Inclusion bodies, containing hnRNPA2 LCD MBP, were pelleted by centrifugation at 24 000 x g for 1h at 4°C. The pellet was resolubilized in a denaturing buffer: 100 mM KCl, 50 mM Tris-HCl, 1 M NaCl, 10 mM imidazole, 1mM dithiothreitol supplemented with 0.1mM PMSF, 0.5 mM BA, and 1 tablet Roche complete EDTA-free protease inhibitor and centrifuged for 1 h at 24 000 x g and 4°C to pellet bacterial debris. The supernatant was filtered through a filter of 0.45 μm and loaded onto a HiPrep 26/10 desalting column using a 20 mM Tris-HCl, 200 mM NaCl, 1 mM DTT, pH 7.4 buffer. Next, the protein samples were combined again, and the solution was loaded onto MBP-trap HP column and eluted with 20 mM tris-HCl, 200 mM NaCl, 1 mM DTT, 10 mM maltose, pH 7.4 buffer.

### Initiating phase separation

#### hnRNPA2 LCD

We observed that the hnRNPA2 LCD is relatively stable in 10 mM CAPS buffer at pH 11.0, but rapidly phase separates when its pH was decreased to 7.5. Thus, we kept the protein in aliquots in a 10 mM CAPS at pH 11.0 and initiated phase separation, first by diluting the protein to the appropriate concentration in the same buffer, then starting the reaction by decreasing the pH with 1.6 μl of 0.5 M MES pH 5.5 for every 100 μl of protein solution to a final value of 7.5. However, for each batch and different protein concentrations a proper amount of MES should be carefully adapted. Phase separation commenced immediately and could be then followed and characterized by a variety of techniques.

#### TDP 43 LCD

TDP 43 LCD is stable in 20 mM MES buffer at pH 3.5 and phase separates when pH increases to 7.5. Therefore, the protein was stored under these conditions and phase separation was induced by applying an appropriate volume of 0.5 M CAPS buffer, pH 11.0 to orangeuced pH up to 7.5. Usually, for 200 μl of TDP 43 LCD (55 μM) we added 5 μl of CAPS buffer (0.5 M, pH 11.0).

#### NUP98-LCD

NUP 98-LCD is stable in 50 mM PBS buffer at pH 3.5 and phase separates when pH increases to 10.0. Therefore, the protein was stored under these conditions and phase separation was induced by applying an appropriate volume of 0.1 M CAPS buffer, pH 11.0 to orangeuced pH up to 10.0.

### Fluorescent labelling with Dylight® 488

#### hnRNPA2 LCD

100 μl of of 8 mg/ml hnRNPA2 LCD in TEV cleavage buffer (3 M urea, 200 mM NaCl, 50 mM NaH_2_PO_4_) was dialyzed against 0.1 M sodiumcarbonate buffer, pH 8.5. 10 mg/ml of the fluorescent Dylight® 488 dye (Thermo scientific) was dissolved in DMSO and added to the protein at a final concentration of 0.05 mg/ml. The solution was incubated at room temperature for 1h and then was dialyzed against 0.01 M CAPS, pH 11.0 storage buffer, to remove the excess of unbound fluorophore. Fluorescently labelled hnRNPA2 LCD was protected from light and stored at −80°C.

#### TDP 43 LCD

100 μM of Dylight® 488 dye (Thermo scientific) was used to label TDP 43 LCD for fluorescent microscopy measurements. TDP 43 LCD (55 μM) in 1x PBS was added to the dye and incubated at room temperature for 30 min in darkness. The mixture was then desalted with the Zeba™ Desalting Spincolumns, 7K MWCO (Thermo Fisher Scientific) to the reaction buffer (1x PBS, 10 mM MgCl_2_, 0.05% Tween 20). Fluorescently labelled TDP 43 LCD was protected from light and stored at −80°C.

### Turbidity measurements

In addition to the MES buffer and/or CAPS buffer to manipulate the pH, appropriate volumes of sodium chloride (Sigma®) was administered to a non-binding black 96 well plate with transparent bottoms. The following protein solutions were used: i) 200 μl of 20 μM hnRNPA2 LCD in CAPS buffer, pH 11 was mixed with 1.6 μl of 0.5 M MES pH 5.5 to reach a final pH equal to 7.5, ii) 200 μl of 55 μM TDP 43 LCD in 20 mM MES, pH 5.5 buffer was mixed with 5 μl of 0.5 M CAPS, pH 11.0 buffer to reach a final pH equal to 7.0. In order to measure phase separation at different concentrations, a volume of MES or CAPS buffer should be carefully adapted. The protein solution was simultaneously added to each well containing a desired buffer, and after a short mixing, the plate was covered with a transparent film (VIEWsealTM) and the absorbance of the solution was measured at 600 nm for 6 hours on a BioTek SynergyTM Mx plate reader at 25°C with continuous shaking. Experiments were conducted in triplicate and then averaged.

### Dynamic light scattering measurements

Dynamic light scattering (DLS) measurements were carried out on a DynaPro NanoStar (Wyatt) instrument. A disposable cuvette (WYATT technology) was filled with 100 μl of protein solution at pH and concentration values at which phase separation was observed. The sides of the cuvette were filled with water and a cap was put on top. The intensity of scattered light was recorded at a scattering angle of 95° at 25°C for a period of 6 hours collecting 10 acquisitions (8 s each). Each measurement was repeated at least 3 times. The software package DYNAMICS 7.1.9 was used to analyse the data, and particle size was calculated based on the following formulas:

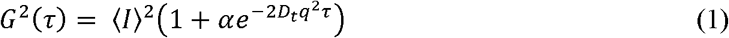

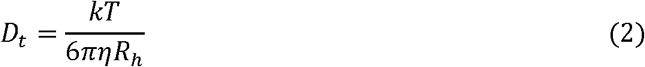

where *G*^2^ (*τ*) is autocorrelation function, *D*_*t*_ is the hydrodynamic radius, *τ* is delay time, *q is* refractive index, *D*_*t*_ is diffusion coefficient, *k* is the Boltzmann’s constant, *T* is temperature, *η* is viscosity of fluid, *R*_*h*_ is hydrodynamic radius.

### Fluorescent and brightfield microscopy

Microscopy measurements were carried out on a Leica DMi8 microscope. Dylight® 488-labeled hnRNPA2 LCD or TDP 43 LCD were mixed with non-labelled hnRNPA2 LCD or TDP 43 LCD (at a 1:200 ratio). Then, phase separation was induced by changing the pH of the protein solution as described earlier. The solution was incubated at 25°C and droplets were visualized with 100 times magnification with brightfield, and/or fluorescence microscopy (FITC filter).

### Statistical analysis

Statistical analysis was performed in Microsoft Office Excel. All data are shown as mean ± standard deviation.

