## Supplementary material for "A generic approach for studying the kinetics of liquid-liquid phase separation under near-native conditions": Figure S1

*Corresponding authors:

**Figure S1. Domain organization of full-length hnRNPA2, TDP43 and NUP98, and amino-acid sequence of their LCD**

Source: http://www.uniprot.org/

a) hnRNPA2-LCD

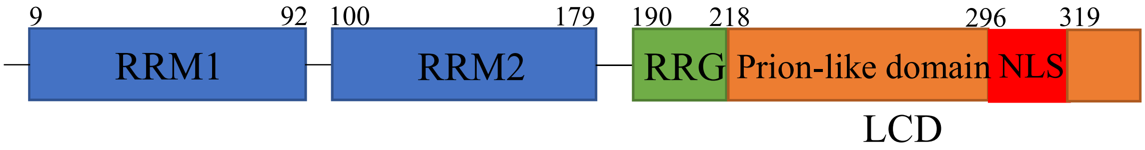


RSGRGGNFGFGDSRGGGGNFGPGPGSNFRGGSDGYGSGRGFGDGYNGYGGGPGGGN
FGGSPGYGGGRGGYGGGGPGYGNQGGGYGGGYDNYGGGNYGSGNYNDFGNYNQQ
PSNYGPMKSGNFGGSRNMGGPYGGGNYGPGGSGGSGGYGGRSRY

Molecular weight: 14607.91
Theoretical pI: 9.45

b) TDP 43- LCD, fragment: 267-414

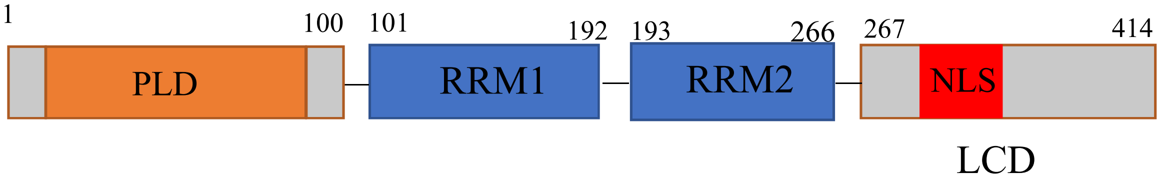


NRQLERSGRFGGNPGGFGNQGGFGNSRGGGAGLGNNQGSNMGGGMNFGAFSINPAMMA
AAQAALQSSWGMMGMLASQQNQSGPSGNNQNQGNMQREPNQAFGSGNNSYSGSNSGA
AIGWGSASNAGSGSGFNGGFGSSMDSKSSGWGM

Molecular Weight: 14564.51

Theoretical pI: 10.75

c) NUP 98- LCD, fragment: 2-299

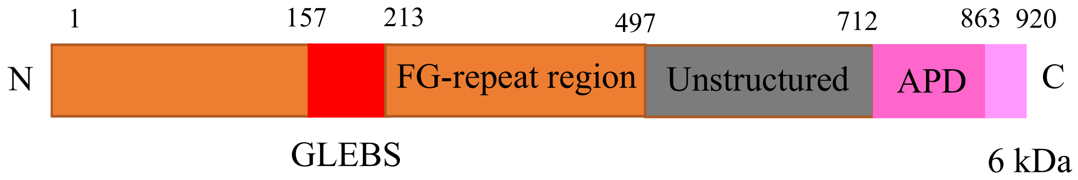


MFNKSFGTPFGGGTGGFGTTSTFGQNTGFGTTSGGAFGTSAFGSSNNTGGLFGNSQTKPGGL
FGTSSFSQPATSTSTGFGFGTSTGTANTLFGTASTGTSLFSSQNNAFAQNKPTGFGNFGTSTSS
GGLFGTTNTTSNPFGSTSGSLFGPSSFTAAPTGTTIKFNPPTGTDTMVKAGVSTNISTKHQCIT
AMKEYESKSLEELRLEDYQANRKGPQNQVGAGTTTGLFGSSPATSSATGLFSSSTTNSGFAY
GQNKTAFGTSTTGFGTNPGGLFGQQNQQTTSLFSKPFGQATTTQNTGFSFGNTSTIGQPSTNT
MGLFGVTQASQPGGLFGTATNTSTGTAFGTGTGLFGQTNTGFGAVGSTLFGNNKLTTFGSST
TSAPSFGTTSGGLFGNKPTLTLGTNTNTSNFGFGTNTSGNSIFGSKPAPGTLGTGLGAGFGTA
LGAGQASLFGNNQPKIGGPLGTGAFGAPGFNTTTATLGFGAPQAPVALTDPNASAAQQ

Molecular Weight: 48971.83

Theoretical pI: 9.81
